# It matters who you know, down below: Different mycorrhizal fungal communities differentially affect plant phenolic defences against herbivory

**DOI:** 10.1101/2020.07.06.190710

**Authors:** Adam Frew, Bree A. L. Wilson

**Affiliations:** School of Sciences, University of Southern Queensland, Toowoomba, QLD, Australia; Centre for Crop Health, University of Southern Queensland, Toowoomba, QLD, Australia; Institute for Land, Water and Society, Charles Sturt University, Wagga Wagga, NSW, Australia

**Keywords:** Herbivory, multitrophic interactions, mutualism, mycorrhizal fungi, plant defence

## Abstract

Arbuscular mycorrhizal (AM) fungi are ubiquitous symbionts of most terrestrial plants. These fungi not only provide their host plants with access to nutrients and resources but are known to augment plant defences against insect herbivores. Relatively little is known about the role of AM fungal diversity and community assembly on the expression of plant defence traits. Here, we report how plant (*Triticum aestivum*) phenolic-based resistance to insect herbivory is differentially affected by different AM fungal communities. An inoculant of four AM fungal species and a field-sourced native AM fungal community increased plant phenolics by 47.9% and 50.2%, respectively, compared to plants inoculated with only one fungal species. Correspondingly, the performance (relative growth rate) of the insect herbivore was 36% and 61.3% lower when feeding on plants associated with these AM fungal communities. Furthermore, there was a negative correlation between foliar phenolics and herbivore growth. We propose that AM fungal community assembly can drive insect herbivore performance by affecting phenolic-based defence mechanisms.

## INTRODUCTION

Most terrestrial plants form a symbiotic relationship with arbuscular mycorrhizal (AM) fungi (Subphylum: Glomeromycotina). These fungi colonise plant roots and provide their host plants with access to important resources such as phosphorus, while the plants provide carbon to the fungi in the form of sugar and lipids [1]. In addition to increasing nutrient acquisition, AM fungi can enhance plant growth and help plants to cope with attack from insect herbivores [2,3].

When challenged with herbivory, plants can rely on different defences to limit damage. Resistance-based defences form a key plant defence strategy utilising chemical or physical mechanisms that will deter insect herbivores or reduce their performance (e.g. survival, growth, fecundity). The AM symbiosis can enhance the capacity for plants to defend themselves. For example, AM fungi can increase plant phenolic compounds [4], silicon concentration [5] and other secondary metabolites associated with resistance to herbivores [3].

Evidence suggests not all AM fungal taxa deliver the same outcome for their plant hosts, and AM fungi vary in their expression of traits associated with different functions (i.e. plant growth, nutrient uptake, defence)[6]. For example, certain taxa have been more associated with augmenting plant nutrient uptake, and others with increasing plant defences against pathogens [7]. Furthermore, the outcome of the AM symbiosis can also depend on the identity of the host plant. For example, recent research has shown plant growth and nutrient status are differentially affected by different AM fungal communities, an effect which also varies between plant species [8].

Moreover, evidence shows that AM fungal species vary in how they affect plant responses to herbivory [9], and insect performance [2]. The AM symbiosis can increase plant phenolic compounds [4] which are multi-functional, but can play an important role in defence against herbivory in many plants [10,11]. Yet, to date we do not know how different AM fungal communities might differentially alter plant phenolic-based resistance to insect herbivory. Therefore, we investigated how different communities of AM fungi affect plant foliar phenolics in wheat (*Triticum aestivum*) and the effects on the performance of a generalist insect herbivore (*Helicoverpa punctigera*).

## METHODS

### Experimental set-up

In a factorial pot experiment, *Triticum aestivum* L. (wheat) cv. Hindmarsh were grown, singly, in 3.7L pots (3.42 kg oven dry equivalent) of gamma-sterilised 80 : 20 soil : quartz sand mixture (Table S1). Plants were inoculated after 7 d with one of four AM fungal (AMF) treatments using approximately 300 AM fungal spores pipetted directly onto roots. All spores were extracted using the wet sieving and sucrose centrifugation method [12]. The treatments were (i) **No AMF** inoculum comprising of equal proportions of sterilised (autoclaved) ‘one AMF species’ ‘four AMF species’ and ‘native AMF’ inoculants; (ii) **One AMF species** using a commercial inoculum (Microbe Smart Pty. Ltd. Melrose Park DC. South Australia) containing only *Rhizophagus irregularis*; (iii) **Four AMF species** using a commercial inoculum containing four species identified as *Claroideoglomus entunicatum, Funneliformis coronatum, Rhizophagus irregularis* and *F. mosseae*; (iv) **Native AMF community** comprising AM fungal spores extracted from the field soil (Table S1). All pots received microbial filtrate (300 ml) to standardise the microbial community within each pot at the initiation of the treatments as per Frew et al. [8].

Plants were grown in a growth chamber (Conviron® PGW40) with day : night temperatures of 27 °C (±4 °C) and 17 °C (±4 °C) respectively, daylight set at 900 mol^-2^s^-1^ with a 12h photoperiod and relative humidity controlled at 60% (±8%). Every two weeks pots were rearranged to reduce any spatial effects. After eight weeks, a fourth-instar *Helicoverpa punctigera* larva was weighed and placed on half of the plants under each AM fungal treatment (10 replicates, nine replicated for the ‘No AMF’ plants, due to the death of one replicate plant). To prevent the larvae escaping, a 45 cm x 55 cm polyester (organza) sleeve was secured around all plants with elastic bands (including insect-free plants). After 6 days, larvae were removed from the plants, and starved for a further 12 h before being reweighed to calculate relative growth rate (RGR) as per [5]. Plants were removed from their pots, the roots and shoots were separated (1g root subsample was randomly removed for mycorrhizal scoring) and the remaining plant material was snap-frozen, freeze-dried, and weighed.

### Mycorrhizal scoring

To measure mycorrhizal colonisation of roots, 1g sample of roots from each plant was cleared with 10% KOH in a 90 °C water bath for 15 mins and then stained with 5% ink-vinegar for a further 10 mins [13]. Stained roots were scored for the presence of AM fungi using the grid intersect method [14] for a minimum of 100 intersects.

### Plant chemical analysis

Freeze-dried leaf material was ground to a fine powder using a plant tissue lyser, and analysed for carbon and nitrogen concentration using a CHN analyser (LECO TruSpec Micro, LECO Corporation, Saint Joseph, MI, USA). Total phenolic concentration was determined as described in Salminen and Karonen [15] as per Frew et al. [16].

### Statistical analysis

R statistical interface (v3.6.1) was used for all statistical analysis [17].

Data exploration for all responses was carried out following the protocol described in Zuur, Ieno, and Elphick (2010).

The effects of the AM fungal treatments and the herbivore treatment on aboveground and belowground biomass, phenolics, C:N and AM fungal root colonisation were assessed by fitting standard linear models using the *lm* function [19] initially comparing factors ‘AM fungi’, ‘herbivore’ and their interactions. Belowground biomass was log transformed to meet model assumptions. There was no evidence of AM fungal colonisation in the ‘No AMF’ treatment and thus were not included in that model. To determine which predictors were important for each of these response variables, the *dredge* function, from the R package ‘MuMIn’ [20], generated combinations of models where the second-order Akaike Information Criterion (AICc) was used to compare the level of support for models within the candidate set [21,22]. Models in which the AICc difference is <2 are considered to have substantial support (Burnham & Anderson, 2003). Where either or both factors were important predictors, effect sizes were calculated (R^2^), ANOVAs from the package ‘car’ [19] and Tukey’s pairwise comparison using the *HSD.test* function ‘agricolae’ [23] were applied to those models with most support (Table 1).

**Table 1.**
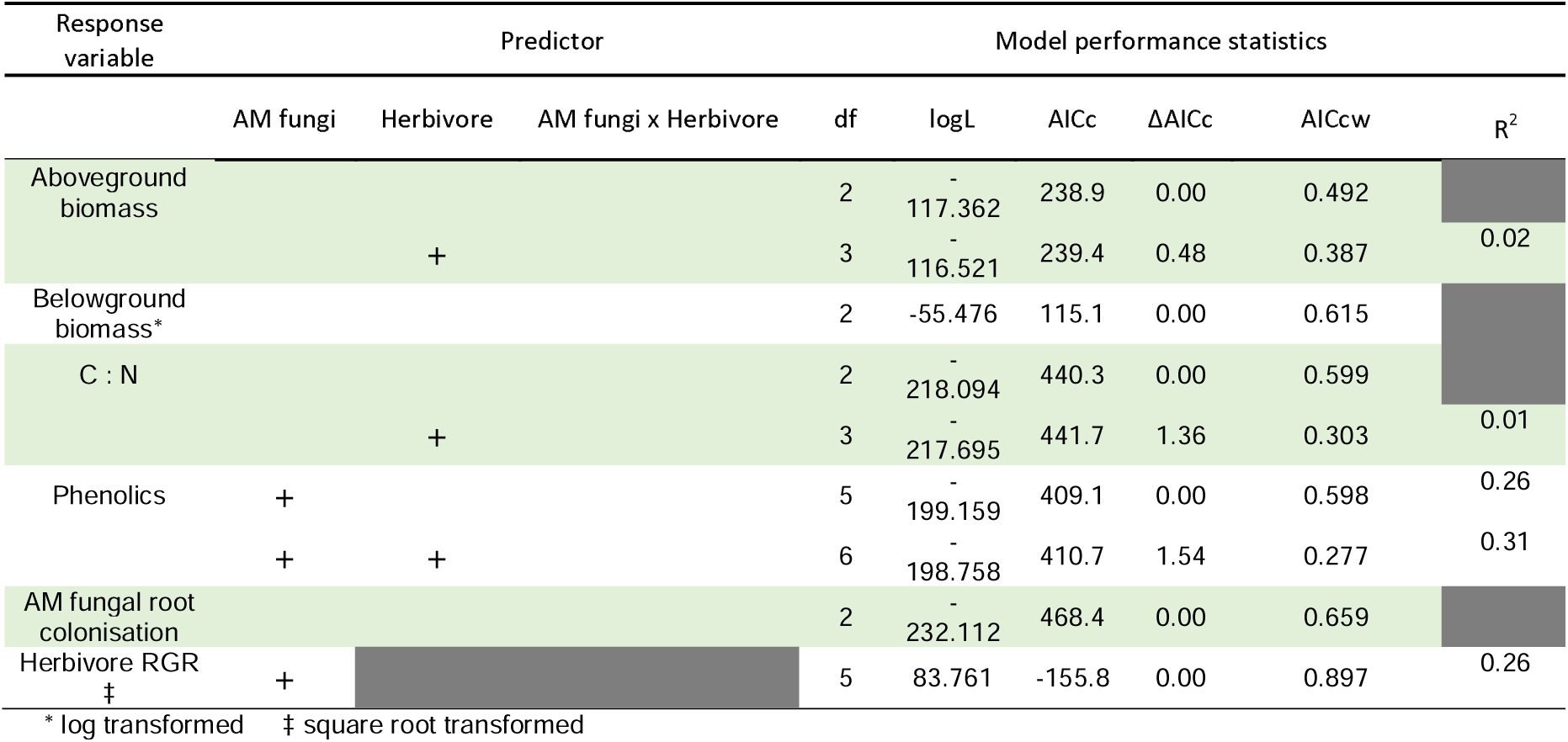
Model selection process for the simple linear models applied in the analysis of the response variables. Predictors included in the models are indicated with crosses. Where the best model has no crosses, no predictor explains variation in response variable. The degrees of freedom (df), log likelihood score (logL), corrected Akaike’s information criterion (AICc), difference in AICc relative to the best model (ΔAICc) and AICc weight (AICcw) are presented for each model. Only models with a ΔAICc ≤ 2 are shown.

The effects of the AM fungal treatments on herbivore relative growth rates was assess using a standard linear model with ‘AM fungi’ as the predictor, RGR was square root transformed to meet model assumptions. Effect sizes were calculated, ANOVAs and Tukey’s pairwise comparison used as before [23]. The relationship between insect herbivore RGR and foliar phenolic concentrations was assessed using standard linear model and Pearson’s correlation coefficient using the *cor.test* function [19].

## RESULTS

Aboveground biomass and belowground biomass were not affected by herbivore or the AM fungal treatments, and neither predictor explained variation in plant biomass (Table 1; Figure S1). Foliar C : N was similarly unaffected by any of the treatments (Table 1; Figure S2). Root colonisation by AM fungi was not different between the three mycorrhizal treatments, and was unaffected by the herbivore treatment (Table 1; Figure S3). No colonisation was detected in the ‘no AMF’ controls.

Model selection indicated that AM fungi were a significant predictor of plant total phenolics (R^2^ = 0.26; Table 1). Post-hoc analysis showed plants inoculated with four AM fungal species and with the native AM fungal community had 47.9% and 50.2% greater phenolics (P<0.001), respectively, than plants inoculated with one AM fungal species (Figure 1a; Tables 1, 2).

**Table 2.** Results of ANOVAs on the simple linear models with the most support (see Table 1). Significant impacts (*P*<0.05) are indicated in bold.

|  | AM fungi |  | Herbivore |  |
| --- | --- | --- | --- | --- |
|  | F | P | F | P |
| Aboveground biomass |  |  | 1.66 | 0.2 |
| C : N |  |  | 0.78 | 0.38 |
| Phenolics | **8.33** | **<0.001** |  |  |
| Herbivore RGR | **4.1** | **0.01** |  |  |

**Figure 1.**
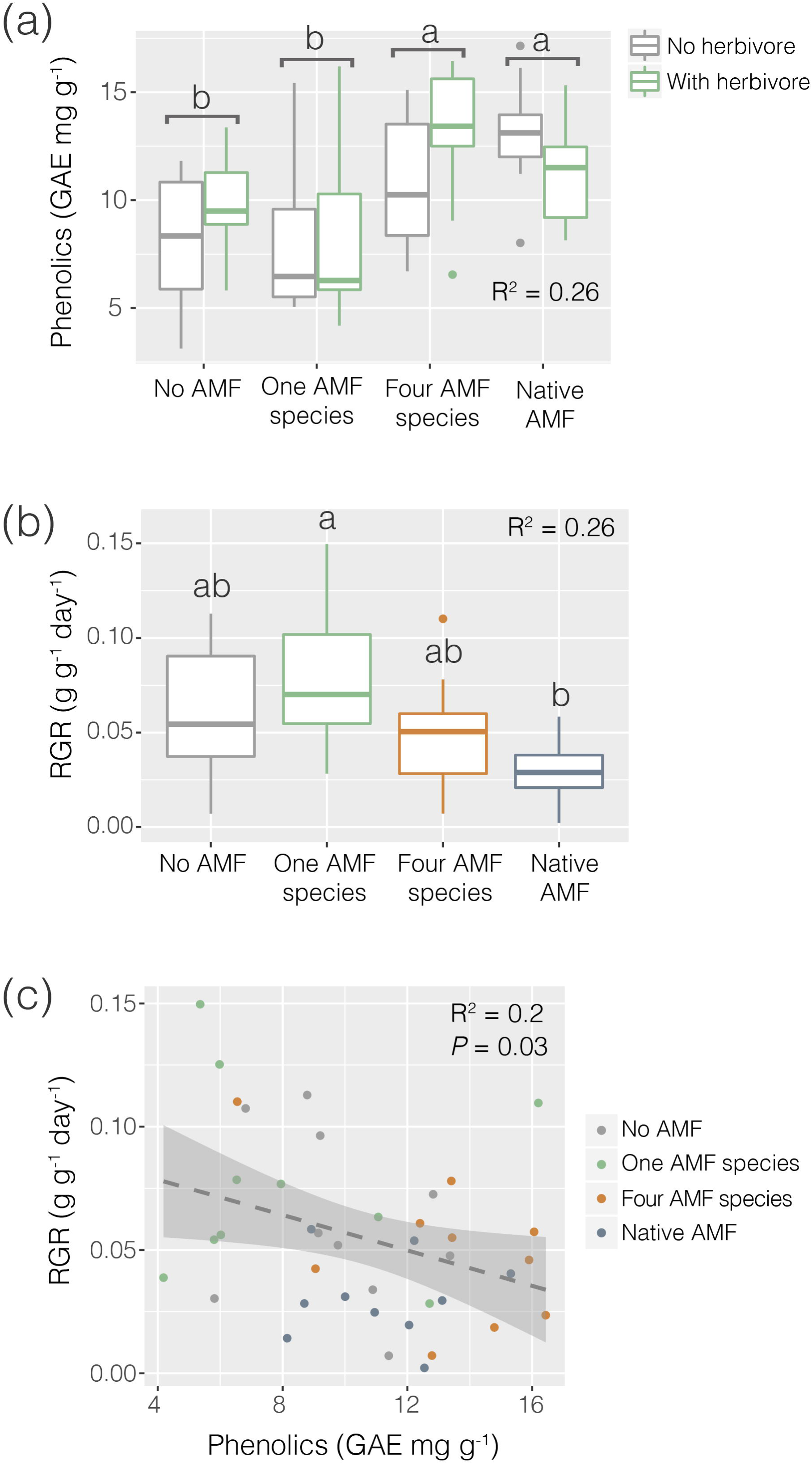
(a) Effects of herbivory on total foliar phenolics (GAE mg g^-1^) of plants (*Triticum aestivum*) grown with no arbuscular mycorrhizal fungi (AMF), a single AMF species, four AMF species or with a native AMF community. **(b)** The relative growth rate (g g^-1^ day^-1^) of a generalist herbivore (*Helicoverpa punctigera*) feeding on plants grown with no AMF, a single AMF species, four AMF species or with a native AMF community. Where factor effects are significant, boxes not sharing a common letter differ significantly (*P*<0.05). **(c)** Relationship between *H. punctigera* relative growth rate (RGR) and foliar phenolics, dashed line represents linear regression through all points and shaded area represents 95% confidence intervals. Coefficients of determination (R^2^) and *P*-value are shown where appropriate.

Model selection indicated that AM fungi were a significant predictor of *H. punctigera* RGR (R^2^ = 0.26; Tables 1, 2). Growth rates were lowest on plants inoculated with the native AM fungal community. This was 51% and 61% lower compared to those herbivores fed on plants inoculated with no AM fungi or with one AM fungal species, respectively (Figure 1b; *P* = 0.01). Post-hoc analysis showed those insects which fed on plants inoculated with the native AM fungal community had significantly lower RGR compared to those inoculated with one AM fungal species. *Helicoverpa punctigera* RGR was negatively correlated with foliar phenolics (Figure 1c; R^2^ = 0.2, *P* = 0.03).

## DISCUSSION

Our study shows different AM fungal communities can differentially affect plant resistance to insect herbivores through changes in foliar phenolics. Plants inoculated with either the native or four AM fungal communities had more phenolics compared to non-mycorrhizal plants, or those inoculated with only *R. irregularus*. Augmented phenolics corresponded to lower growth rates of the generalist insect herbivore feeding on those plants.

It is well-established that outcomes of the AM symbiosis for the host plant can vary depending on AM fungal identity. This has been shown for plant defences to pathogens [7] and responses to herbivory [9,24] but has, until now, not been associated to particular defence mechanisms. In this instance, inoculation with particular AM fungal communities increased foliar phenolics where insects on these plants had the lowest performance of all the treatments. This finding was supported by the negative correlation between phenolic concentration and insect growth.

Although plants under the single AM fungal species treatment were well colonised, consisting only of *R. irregularis*, there was no effect on plant biomass, nutritional quality (C:N), or phenolics, nor did it affect herbivore performance. As a cosmopolitan and generalist species, this itself is noteworthy. Although there is notable variability in both plant [8] and insect herbivore responses to this fungal species in the literature [2,25]. In our study, the inclusion of three additional AM fungal species along with *R. irregularis* resulted in phenolic-based herbivore resistance, with similar outcomes for plants inoculated with the native AM fungal community. Thus, increasing AM fungal species richness (although the native fungal community was not identified) potentially increased opportunity to engage in mycorrhizal associations with fungal species that can promote plant defence. Therefore, the four AM fungal species and the native community may include a particular species that enhances plant phenolics, or it may be that a combination of functionally complementary AM fungi together resulted in elevated phenolics in the leaves of their shared host plant. Although our understanding of the role of AM fungal diversity in plant defence against herbivores is in its infancy, there is growing evidence of functional complementarity among AM fungi, at least with regard to plant pathogen protection [26].

Despite increasing emphasis on the community approach to studying fungal-mediated above-belowground interactions [27], empirical data on how AM fungal community assemblage affects the expression of plant defence traits is still limited. The distinct defence outcomes conferred by different AM fungal communities here suggests potential for selection towards associations with particular AM fungal taxa, especially in plant communities subject to greater herbivore pressure. Furthermore, some plant taxa may have a competitive advantage over others based on their evolved defence mechanisms. For example, in soil environments where the four AM fungal species used in our study are present, plants with evolved capacity and dependence on phenolic-based defences in this case could be more competitive than plant taxa reliant on other evolved defence mechanisms. Such a competitive advantage would more than likely to be short-lived however, as the derived ‘benefit’ of engaging in the AM symbiosis is environmentally context dependent (e.g. soil fertility). Thus, the stability of these relationships in natural systems would occur only in certain circumstances. For managed plant production systems, particularly environments of intensive agricultural activity, our results highlight the potential for the application of tailored AM fungal inoculants suited to the local environment, for particular crop species. Our findings also highlight that in some instances, the application of AM fungal inoculants may provide no additional benefit than the native fungal community already present in the soil, as we found here, perhaps due to functional redundancy of additional AMF taxa.

Our study demonstrates that different AM fungal communities can differentially affect phenolic-mediated plant resistance to an insect herbivore, highlighting that AM fungal species identities are an important consideration for herbivore protection. Our findings also suggest that the assembly of resident AM fungi in some contexts may offer effective mycorrhizal-induced herbivore protection, and thus land management practices that favour AM fungi could be implemented [28,29]. Nevertheless, considering the highly context-dependent outcomes of the AM-symbiosis more studies are necessary to build on our findings and assess these effects across a range of AMF communities, plant hosts, and environmental contexts, as the outcomes are sure to vary.

## DATA ACCESSIBILITY

Raw data are available from the Dryad data Repository: DOI to be supplied upon acceptance.

## ACKNOWLEDGEMENTS

The authors would like to thank the technical team at Charles Sturt University for their technical support for this project. The authors would also like to thank Dale Nimmo for his advice on statistical analyses.

## AUTHOR CONTRIBUTIONS

A.F. obtained funding, conceived and designed the study, and acquired the experimental data. Data were analysed by A.F. with input from B.A.L.W.. Both A.F. and B.A.L.W. interpreted the data and drafted the manuscript. Both authors approve the final version to be published.

