## Supplementary material for "It matters who you know, down below: Different mycorrhizal fungal communities differentially affect plant phenolic defences against herbivory": Table S1

**Table S1**. Nutrient analysis of fully homogenised soil substrate gamma irradiated (50 KGY). Soil was collected from previously grazed paddock site (142 m a.s.l. elevation) in NSW, Australia (35°02'45.6"S 147°20'53.8"E). Analysis carried out by Environmental Analysis Laboratory, Southern Cross University, Lismore, Australia.

| **Nutrient (Method)** | **Units** | **Soil** |
| --- | --- | --- |
| pH (water) | pH unit | 5.14 ± 0.30 |
| Exchangeable magnesium (ammonium acetate) | mg/kg | 179.5 ± 1.71 |
| Exchangeable calcium (ammonium acetate) | mg/kg | 785.75 ± 13.97 |
| Exchangeable potassium (ammonium acetate) | mg/kg | 304.25 ± 13.29 |
| Exchangeable sodium (ammonium acetate) | mg/kg | 26.5 ± 1.5 |
| Ammonium (KCl) | mg/kg | 12.28 ± 2.43 |
| Total nitrogen (LECO analyser) | % | 0.05 ± 0.01 |
| Total carbon (LECO analyser) | % | 0.56 ± 0.12 |
| Total phosphorus (acid extractable) | mg/kg | 254.75 ± 5.07 |
| Phosphorus (colwell) | mg/kg P | 32 ± 3.34 |
| Phosphorus (Bray1) | mg/kg P | 13.03 ± 2.92 |
