## Supplementary material for "It matters who you know, down below: Different mycorrhizal fungal communities differentially affect plant phenolic defences against herbivory": Figure S1

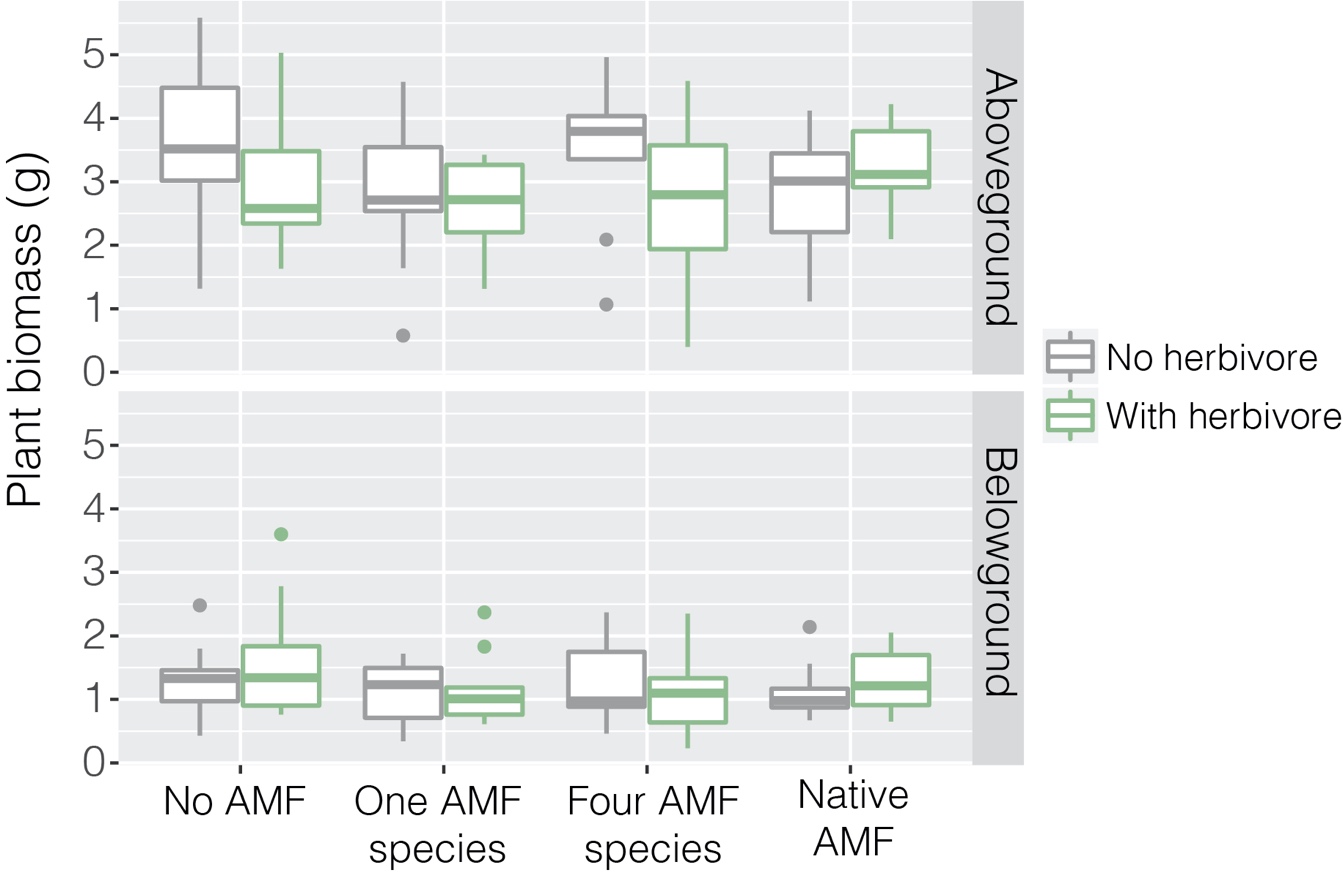

**Figure S1** Effects of herbivory (*Helicoverpa puntigera*) on the aboveground and belowground biomass of wheat (*Triticum aestivum*) plants inoculated with no arbuscular mycorrhizal fungi (AMF), a single AMF species, four AMF species, or a native AMF community.
