## Supplementary material for "It matters who you know, down below: Different mycorrhizal fungal communities differentially affect plant phenolic defences against herbivory": Figure S3

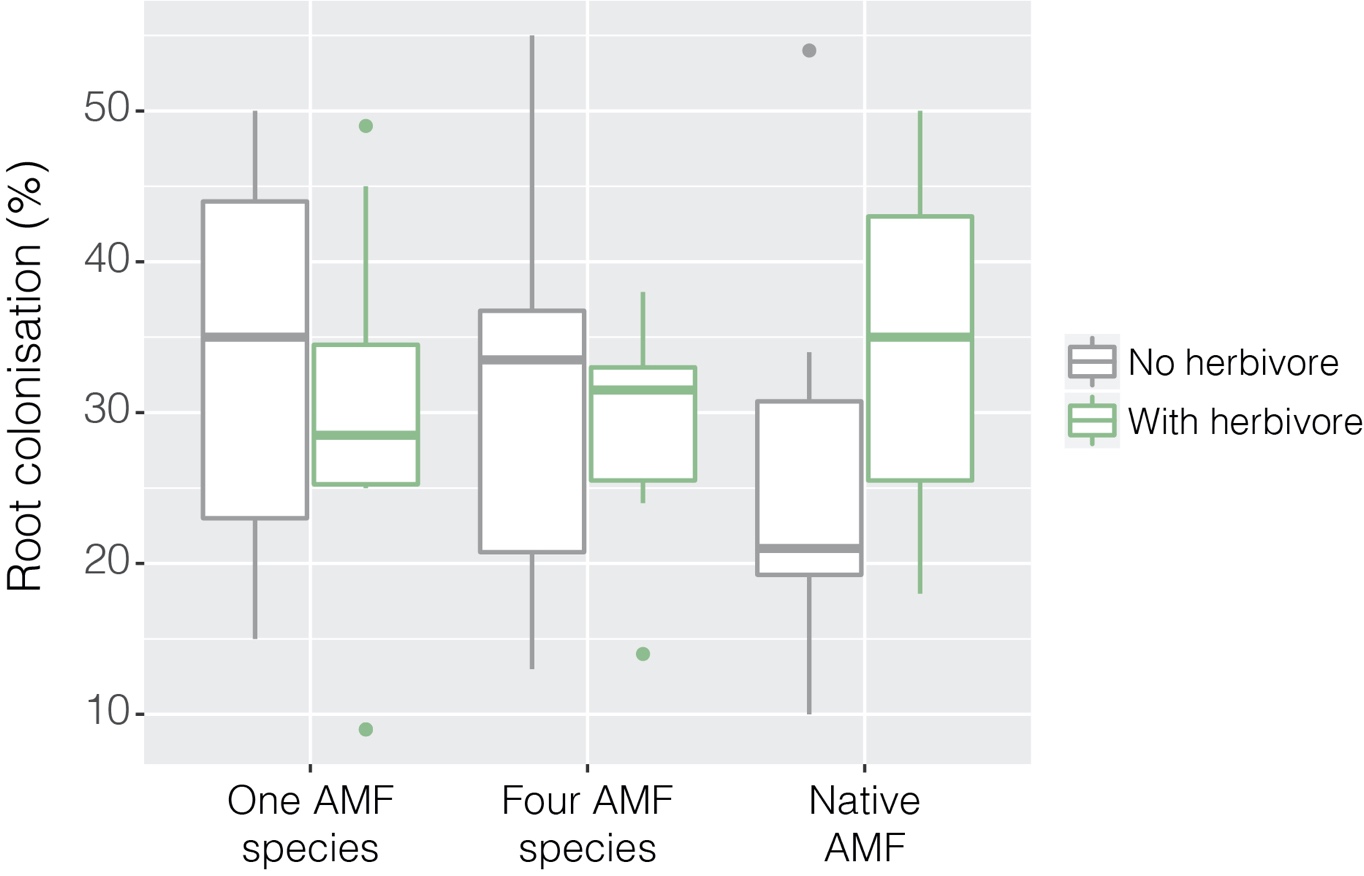

**Figure S2** Effects of herbivory (*Helicoverpa puntigera*) on the total arbuscular mycorrhizal fungal (AMF) root colonisation of wheat (*Triticum aestivum*) plants inoculated with no AMF, a single AMF species, four AMF species, or a native AMF community.
